# Knowledge-based classification of fine-grained immune cell types in single-cell RNA-Seq data with ImmClassifier

**DOI:** 10.1101/2020.03.23.002758

**Authors:** Xuan Liu, Sara J.C. Gosline, Lance T. Pflieger, Pierre Wallet, Archana Iyer, Justin Guinney, Andrea H. Bild, Jeffrey T. Chang

## Abstract

Single-cell RNA sequencing is an emerging strategy for characterizing the immune cell population in diverse environments including blood, tumor or healthy tissues. While this has traditionally been done with flow or mass cytometry targeting protein expression, scRNA-Seq has several established and potential advantages in that it can profile immune cells and non-immune cells (e.g. cancer cells) in the same sample, identify cell types that lack precise markers for flow cytometry, or identify a potentially larger number of immune cell types and activation states than is achievable in a single flow assay. However, scRNA-Seq is currently limited due to the need to identify the types of each immune cell from its transcriptional profile, which is not only time-consuming but also requires a significant knowledge of immunology. While recently developed algorithms accurately annotate coarse cell types (e.g. T cells vs macrophages), making fine distinctions has turned out to be a difficult challenge. To address this, we developed a machine learning classifier called ImmClassifier that leverages a hierarchical ontology of cell type. We demonstrate that ImmClassifier outperforms other tools (+20% recall, +14% precision) in distinguishing fine-grained cell types (e.g. CD8+ effector memory T cells) with comparable performance on coarse ones. Thus, ImmClassifier can be used to explore more deeply the heterogeneity of the immune system in scRNA-Seq experiments.

## Introduction

Single-cell RNA-Seq (scRNA-Seq) has emerged as a powerful technique to catalog cell types [1, 2], including immune cells, which play critical roles in a wide range of diseases. In cancer, they have been shown to impact survival, drug resistance and evolution [3, 4]. However, annotation of the cells based on their transcriptional profiles remains a challenge [5, 6] due both to the diversity of cell types as well as ambiguous divisions along the developmental lineage and activation states [7]. That is, while myeloid and lymphoid cells have drastically different transcriptional profiles and are trivial to distinguish, the differences within each lineage are more subtle—specific types of T cells are difficult to identify [8].

Currently, the most commonly used approach to annotate cell types is to start with an unsupervised clustering algorithm (like t-SNE [9] or UMAP [10]) to group cells with similar profiles, and then to manually inspect each cluster for the expression of marker genes that distinguish specific cell types [11]. While conceptually straightforward, using these markers is challenging in practice due to poor expression or dropout [12, 13], low conservation of markers across studies [14], ambiguity of markers [15], lack of reliable markers [16] and transcriptional similarity of cell types [17]. Further, cell type annotations are not yet easily transferred between different datasets, and therefore, each data set needs to be manually annotated by experts with an understanding of both immunology and the idiosyncrasies of scRNA-Seq data. As a final complication, cell type annotation is an iterative procedure where the clustering influences the immune cell classifications, which then reveals discrepancies (e.g. a memory T cell cluster that contains both CD4+ and CD8+ T cells) that need to be resolved by refining the clusters (e.g. by clustering with different genes) [18].

To simplify and automate the process of identifying cell types, several bioinformatics methods have recently been developed. Correlation-based methods such as scmap [19] and SingleR [20] correlate the query cells to a pre-defined set of reference cell types and assign the label of the type with maximum correlation. Hierarchy-based methods such as Garnett [21] and CHETAH [22] construct a reference cell type hierarchy and search for the optimal cell type from a generic root node to increasingly more specific types. CellAssign [23] utilizes a Bayesian statistical framework to model cell types, marker gene expression, and other covariates such as processing batches. While these methods have been successful and quickly adopted into scRNA-Seq pipelines, approaches with improved ability in identifying fine-grained cell types are still needed.

To increase prediction accuracy, and in particular, on more fine-grained cell types, we have developed a method that includes an explicit knowledge-based and hierarchical model of immune cell types. The immune cell types are naturally organized into class hierarchy. ImmClassifier (Immune cell classifier) integrates the biology of immune cells from a hierarchical ontology, and synthesizes heterogeneous reference datasets using a two-step machine learning and deep learning process. Within each reference dataset, the first step trains a random forest classifier to assign probabilities according to the cell types of the reference dataset, which preserves the intra-dataset cell type relations and avoids batch effects when pooling the cells from different reference datasets. To resolve the differences in reference annotations, cell types from all reference datasets were mapped to unified and non-redundant cell ontology hierarchy. The second step employs a deep learning approach to integrate numerous reference datasets and directly learn the cell ontology hierarchy, assigning the optimal annotation based on the distribution of probabilities across the hierarchy. This enables ImmClassifier to synthesize cell type assignments across reference datasets. Evaluating on a number of independent scRNA-Seq datasets, ImmClassifier accurately classified and outperformed existing methods over a variety of immune cells collected from different tissues, especially at coarser annotation granularity. ImmClassifier is available as a Docker container at https://github.com/xliu-uth/ImmClassifier.

## Results

### Evaluating ImmClassifier on tumor-infiltrating immune cells

We developed a computational tool ImmClassifier to annotate immune cells in gene expression data (Methods, Fig 1). We evaluated the performance of ImmClassifier on three test datasets (not included in the training) [17, 24, 25]. We began by analyzing the microarray profiles of purified cell populations, which lacks some of the technical challenges, such as drop-out, that would be seen in the scRNA-Seq data. In this situation, ImmClassifier recovered 13 out of 15 original cell types and distinguished CD4 from CD8 T cells accurately, recovering 100% of CD4 T cells and 83% of CD8 T cells. 95% of the cells identified under the CD4 T hierarchy were CD4 T cells in the original annotation (Fig. 2a). A more challenging situation was seen in the myeloid lineage, where there was less overall coverage in the reference data sets. None of the five eosinophil samples were identified exactly as eosinophils, although three were correctly placed in the myeloid lineage (macrophage, monocytes, dendritic cells). It is likely that greater coverage of this cell type will be needed to improve its accuracy.

**Figure 1.**
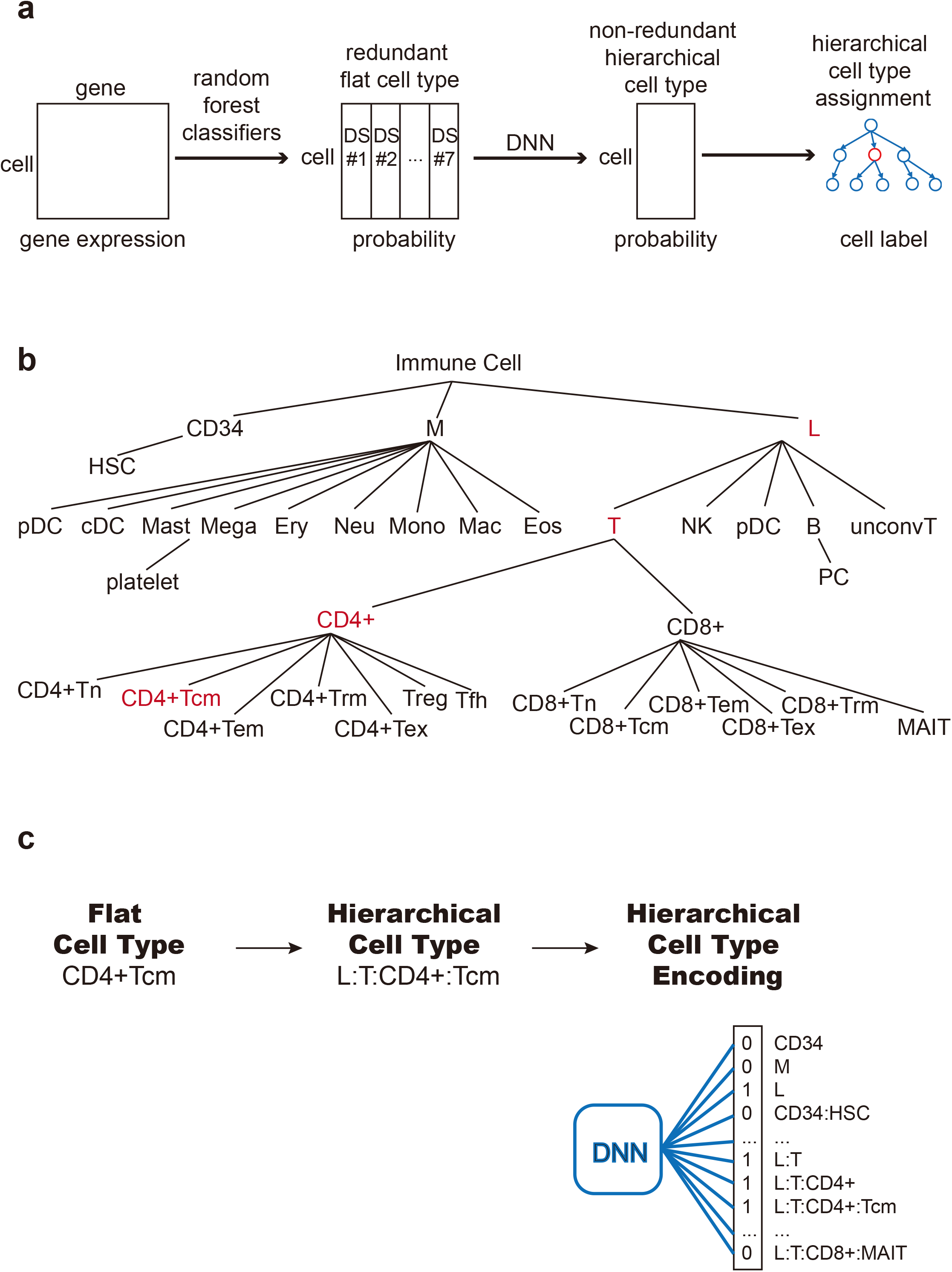
ImmClassifier Architecture. ImmClassifier takes a matrix containing the gene expression of immune cells from a single-cell RNA-Seq experiment (far left). Using feature genes pre-calculated from multiple training datasets, ImmClassifier predicts the probability that each cell corresponds to a cell type annotated in the original training data sets using random forest models (mid left). Here, “DS #1” refers to the first training data set. This probability matrix is further converted to a hierarchical cell type probability matrix using deep neural network classifiers (DNNs). The mean and standard deviation of probabilities from the DNN models is incorporated into a cell ontology hierarchy. Traversing the cell ontology hierarchy, the cell type with maximum information gain is assigned. **(b)** Depiction of the cell types derived from the EBI Cell Onotology to enable machine learning probabilities. Types of immune cells are represented according to granularity, from coarse cell types at the root, to fine-grained ones in the leaf nodes. There are 35 cell types in all. **(c)** Each flat cell type is converted to hierarchical cell type based on the path from root to the flat cell type. The hierarchical cell type is encoded in a 35-bit binary vector by marking the flat cell types on the path 1 and 0 otherwise. The binary vector is used as a target in the DNN training and output in prediction.

To assess its ability to classify scRNA-Seq data, we ran ImmClassifier on tumorinfiltrating immune cells from two distinct cancer types. ImmClassifier achieved an overall precision of 72% and 77% (Fig. 2b, 2c). Again, the highest rate of misclassification was made across closely related lineages. 55% of the B cells were predicted to be plasma cells, which are terminally differentiated B cells (Fig. 2b), which highlight the challenges in distinguishing related closely cell types.

**Figure 2.**
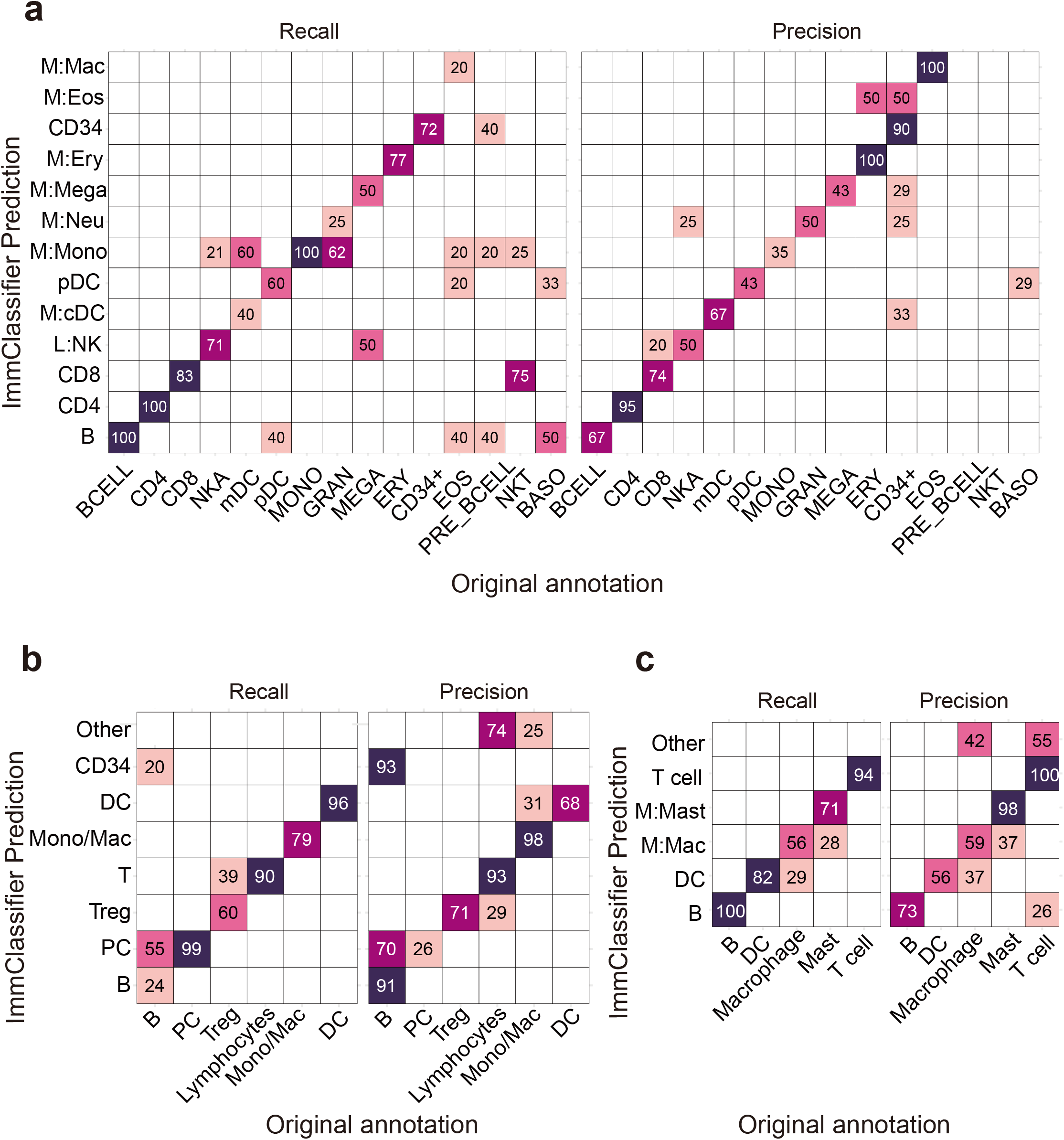
ImmClassifier predicts immune cell types. These heatmaps compare the cell types from the original publication (rows) to those inferred by ImmClassifier (columns). In the recall plot (left), the color represents the recall (as a percent) of each original cell type predicted by ImmClassifier. In the precision plot (right), the color represents the precision of each original cell type predicted by ImmClassifier. Recall or precision scores no less than 20 are labeled. Three datasets tested are **(a)** purified immune populations sequenced by microarray platform [17]. (**b**) SKCM [24]. The *Other* row includes a small number of Megakaryocytes, Mast cells, Eosinophils and Neutrophils. (**c**) HNSCC [25]. The *Other* row includes a minimal number of NK cells, Erythrocytes, Megakaryocytes, Monocytes and Neutrophils. Since ImmClassifier has finer annotation granularity than the original annotation, for the purpose of comparison, the annotation terms of equivalent granularity to the original annotation were used.

### Comparison to existing methods

To determine whether the performance we observe is comparable to related methods, we compared ImmClassifier against Garnett (extended mode) [21] and SingleR [20], which are commonly used representatives for the hierarchy- and correlation-based methods. We applied each method to four scRNA-Seq datasets from three independent publications and visualized the predictions using UMAP clustering. These four datasets cover a variety of sequencing specifications including InDrop 3’ sequencing, 10X 5’ sequencing, 10X 3’ sequencing, Smart-Seq2 Full-length (Table S1). In addition, those datasets covered a wide range of immune populations, collected from cancer patients and healthy donors.

We first quantified the performance of those methods by recall and precision for four different depths across our cell hierarchy (Fig. 3a). (Greater depth is finer cell types). Across all conditions, recall and precision decrease with increasing depth, reflecting the difficulty in distinguishing closely related cell types. For coarse cell types (e.g. myeloid vs lymphoid cells) (depth 1), all methods performed well, with over 95% performance overall. However, the performance dropped with finer types (depth 4) (e.g. central vs effector memory T cells). ImmClassifier achieved an average precision of 97%, 76%, 68%, and 27% for depths 1-4, respectively. At depths 1-2, the recall was comparable to that of SingleR, with only a +3% difference (Fig. 3b). However, at depths 3-4, the recall was improved by +20%. For difference in precision was similar, with a −1% difference at depths 1-2, and +14% at depths 3-4. Thus, while all methods performed well with coarse cell types, ImmClassifier was considerably more accurate at higher depths, although challenges still remain.

**Figure 3.**
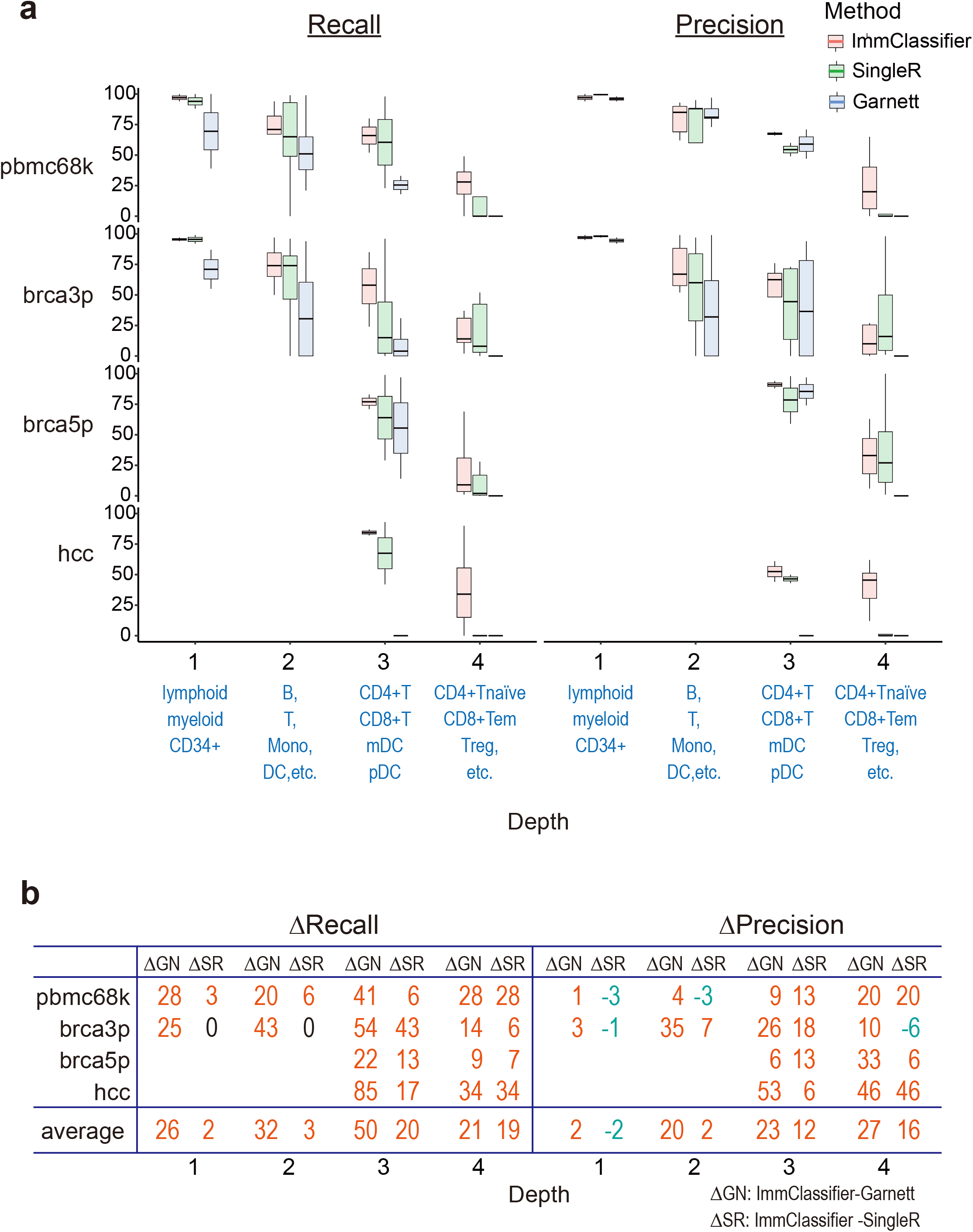
Classification accuracy at different annotation granularities. **(a)** These boxplots show the recall and precision across cell types, organized by depth. Depth 1 includes CD34+, M and L; depth 2 includes HSC, Dendritic, Macrophage, Monocyte, Neutrophil, Mast, T, NK, B; depth 3 includes CD4+T, CD8+ T, mDC, pDC and depth 4 includes CD4+Tn, CD4+Tcm, CD4+Tem, CD4+Tex, CD4+ Treg, CD4+ Tfh, CD8+Tn, CD8+Tcm, CD8+Tem, CD8+Tex and CD8+ MAIT. **(b)** This table shows the average of performance ImmClassifier compared to Garnett and SingleR. The median value of recall/precision across different cell types at each depth for each method was calculated. The difference of median recall/precision (ΔSR, ΔGN) between ImmClassifier and Garnett and SingleR, respectively, was calculated. Comparisons in which ImmClassifier has higher accuracy are shown in red, and those with lower accuracy are blue.

We also compared the methods with respect to (1) spatial concordance with original annotation, (2) frequency of matched and mis-matched cell types to the original annotation, and (3) the distance between the centroids of the predicted and original cell types. We used a data set comprised of a complex mixture of 12 immune cell types, including lymphoid and myeloid cells collected from four tissues [7]. Here, ImmClassifier achieved a higher mean recall and precision across all cell types (45.5% and 43.2%) than SingleR (35.5% and 37.9%) and Garnett (12.5% and 10.7%), with high visual concordance to original annotations in a UMAP plot (Fig. 4a). To assess the overall similarity of the expression profiles of each predicted cell type, we averaged the expression profiles of all cells of each type and computed the Euclidean distance between the averaged profiles with those of the original cells (Fig. 4b). This revealed that the overall profiles of the cell types from ImmClassifier was most similar to those of the original data type. Furthermore, ImmClassifier could recover 11 of the 12 cell types, in contrast to SingleR (8 cell types) and Garnett (6 cell types) (Fig. 4c, Fig.S1). Notably, ImmClassifier was able to distinguish a mast cell population (73% recall, 88% precision), which was completely missed by the other methods (Fig. 4a, 4c). However, all methods failed to identify the cluster of NKT cells. This cell type was missing in the ImmClassifier training set, and both ImmClassifier and SingleR annotated them as a mix of NK cells and T cells (Fig. 4c). Nevertheless, this demonstrates ImmClassifier’s capacity to accurately recapitulate the complexity of the immune cell types.

**Figure 4.**
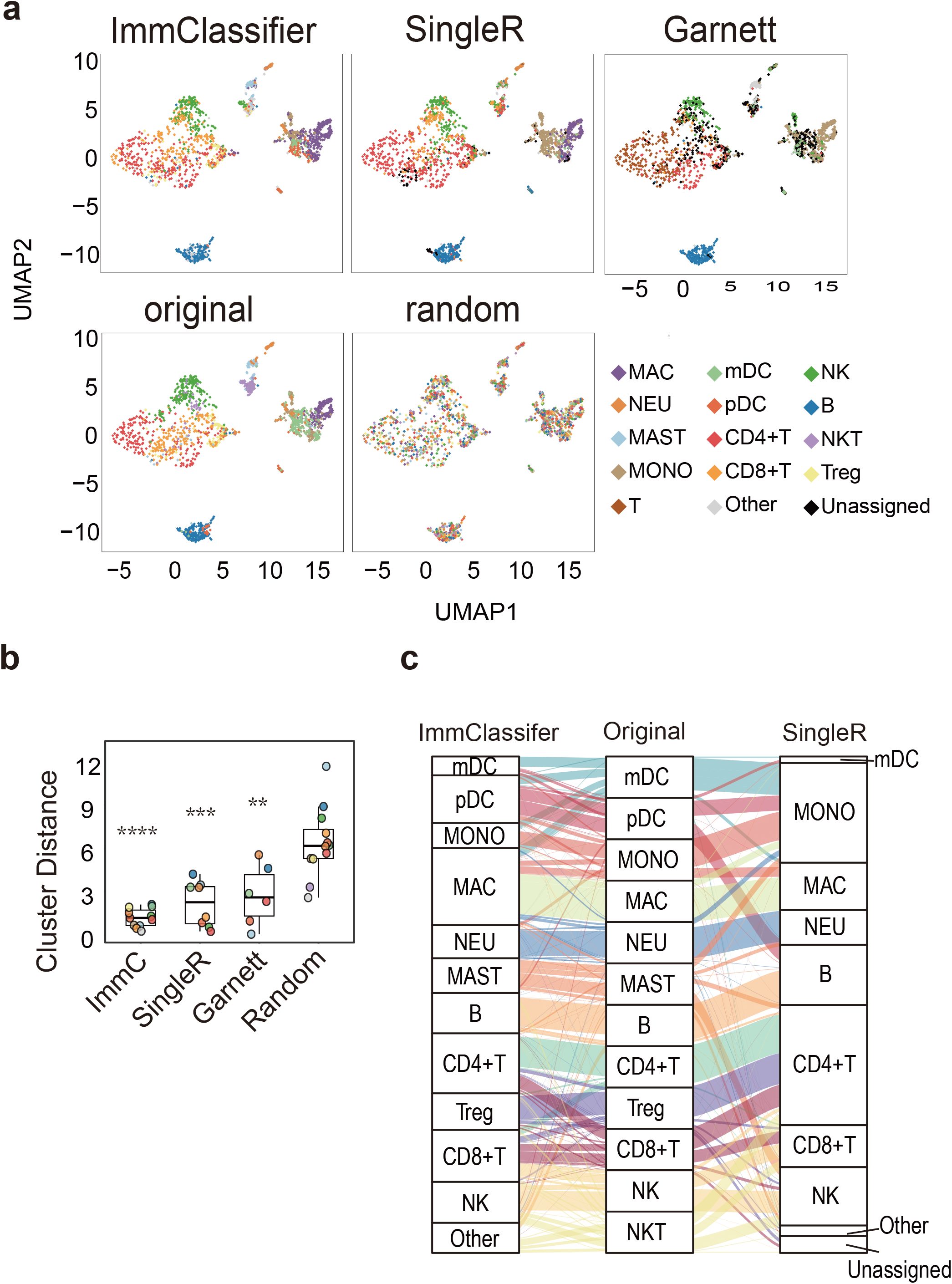
Visualization of brca3p dataset by different annotation methods. For visualization, abundant cell populations (cell number >200) were sub-sampled to 200 per cell type. The mapping IDs of cell types between different annotations is listed in Table S2. If the annotation method produces a finer annotation than the original one, the annotation terms of equivalent granularity were used. **(a)** UMAP plots colored by annotation method. In the first row (left to right), the panels are colored by ImmClassifier, SingleR and Garnett. In the second row (left to right), panels are colored based on the annotations in the original publication, or after random shuffling. **(b)** Boxplots of the Euclidean distance between centroids of predicted annotations and original annotations. P-values are by Wilcoxon test. The significance of P-values is shown as ** (<0.01), *** (<0.001) and **** (<0.0001). (**c**) Alluvial plots connect the original annotated cell types to cell types annotated by ImmClassifier (left) and SingleR (right).

At the deepest level (e.g. distinguishing between central memory and effector memory T cells), the accuracy for ImmClassifier drops to 21.2% recall and 27.1% precision. At this level, the differences in the transcriptional profiles of these cells are subtle, and biological distinctions ambiguous and under heavy investigation [8]. We noticed that ImmClassifier frequently predicted effector memory T cells to be other types of T cells, and in particular, tissue-resident memory (TRM) T cells (Fig. 5a). TRM is a newly identified type of memory T cell that resides in tissue without recirculating, yet is transcriptionally, phenotypically and functionally distinct from recirculating central and memory T cells [26, 27]. It has been shown that TRM cells exhibit high expression of *CXCR6* and low expression *SELL* [28]. To determine whether these cells may be misannotated in the gold standard data set, we examined the ratio of *CXCR6* to *SELL* expression, and see that it is higher in the cells that ImmClassifier predicted to be TRM (Fig. 5b). This suggests that the TRMs were not accurately annotated in the original data set. It is likely that, due to the difficulties in manual annotation, as well as the high degree of expertise required, immune cell subtypes, especially those deep in the hierarchy, are not yet accurately annotated in the existing data sets.

**Figure 5.**
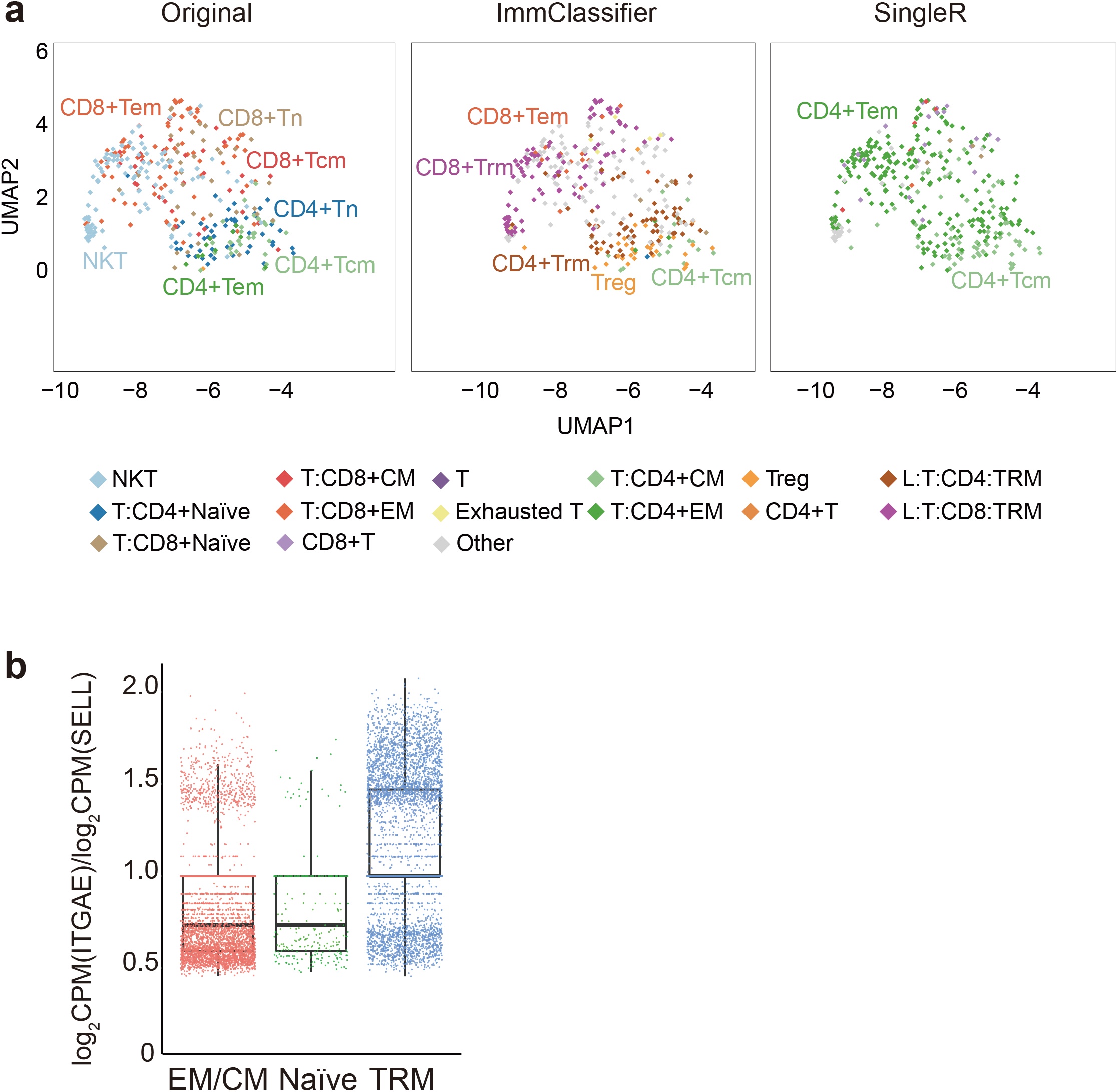
ImmClassifier infers tissue-resident memory T cells in brca5p dataset. **(a)** This shows the T cell clusters from Fig. S1a in greater detail. The UMAP plots are colored by original annotation, ImmClassifier and SingleR. **(b)** This boxplot shows the ratio of the gene expression of TRM marker genes in three cell populations. The *EM/CM* (effector-memory and central memory T) cells are predicted by ImmClassifier, and consistent with the original annotation. The *Naïve* (naïve T) cells are predicted by ImmClassifier and consistent with the original annotation. The *TRM* (effector-memory and central memory T) cells are predicted by ImmClassifier, and inconsistent with the original annotation.

## Discussion

Despite the need for improved annotations of immune cells in scRNA-Seq data, it remains a challenging problem, in particular for cells closely related in the developmental lineage. To address this challenge, the use of a cell type hierarchy has emerged as a critical component in the latest cell type annotation tools. For example, while Garnett is hierarchical, it uses a pairwise classification strategy that does not consider information across the overall ontology, which has been shown in other contexts to improve accuracy [29, 30]. This is in contrast to ImmClassifier, which uses a deep learning framework to model the whole hierarchy and assign the probabilities considering information across all cell types simultaneously, which appears to improve the performance at deeper levels of granularity.

Currently, ImmClassifier cannot detect new and intermediate cell types. ImmClassifier is trained on known cell types and will assign a cell of an unobserved type to its closest known reference cell type. The limit will need to be addressed in future work. To allow identification of intermediate cell types, the continuous distribution of probabilities that ImmClassifier generates in its intermediate step may be informative. But perhaps most importantly, as well annotated scRNA-Seq data sets grow, new reference cell types may be integrated in a straightforward manner within the architecture of ImmClassifier.

We anticipate that the performance of ImmClassifier will continue to increase as additional high-quality data sets become available. Indeed, our results demonstrate the need for a greater quantity and quality of annotated immune cell data set, especially with fine-grained cell types, to train this and other classifiers. The ImmClassifier framework is scalable to new reference data sets, with the most difficult step in mapping the original annotations to the cell hierarchy. We anticipate that ImmClassifier will be most useful in experimental settings that require accurate, comprehensive and robust immune cell annotation.

## Acknowledgement

Research reported in this publication is supported by the National Cancer Institute of the National Institutes of Health under grant number U54CA209978 and the Cancer Prevention and Research Institute of Texas (RP170668). The content is solely the responsibility of the authors and does not necessarily represent the official views of the National Institutes of Health.

## Methods

### ImmClassifier

An outline of ImmClassifier is shown in (Fig. 1a). ImmClassifier employs a machine learning paradigm that takes as input a vector of the expression of *n* genes of a cell and returns an *m* length vector of the probabilities that the cell is from one of the *m* cell types. However, it also deviates from a classical machine learning setup to handle the complexity of the immune cell types and resultant idiosyncrasies in the training sets. Because no single training set includes all desired cell types, multiple ones must be integrated. Furthermore, the cell type labels are frequently inconsistent, using not only different labels, but also labeling at different granularities of the cell type hierarchy. For example, [1] classified T cells into αβ and γδ T cells, [2–4] classified T cells as CD4+ and CD8+ T cells and [5] labeled T cells by numeric cluster ID, without an explicit cell type. To resolve these differences, ImmClassifier takes a stepwise approach where independent classifiers are developed for each training set, and the outputs for each training set are resolved by a final classifier that determines the ultimate cell type assignment. Specifically, the input goes through a random forest classifier for each training set (currently, seven). The output matrices are concatenated and processed by ten independently trained deep neural network classifiers so that a robust estimate of the average performance can be estimated. The mean and standard deviation of the scores for each query cell in each cell type were calculated and projected to a cell ontology hierarchy. The cell type with maximum information gain, relative to its parent node, was assigned to each query cell.

### Reference data sets

To address the heterogeneity of immune cell types and sequencing platforms, seven independent scRNA-Seq datasets covering a wide range of cell types, sources, and sequencing platforms, were used for training (Table S3). Seven independent test data sets were used. Each reference dataset was divided into a training and a test set. The reference datasets are normalized to logarithm of counts per million reads (log_2_ CPM). Non-immune cells were excluded.

### Dataset-specific classification

Feature genes are obtained for each reference dataset. If the cluster-associated marker genes are provided by the original publication, the top 20 marker genes are used. Otherwise, the cell type markers are called by Seurat [6] using the gene expression count and provided cell type annotations. To account for drop-out in scRNA-Seq data, only genes expressed in the query dataset are used.

To reduce the bias introduced by cluster size, a balanced training set is generated by randomly selecting 500 cells per cell type (without replacement for abundant cell types (>1000 cell) and with replacement for rare cell types (<1000 cell)). Batch correction is performed between the query dataset and a reference dataset on feature genes using ComBat [7]. The original cell type labels from the reference dataset were used as the target variable. A random forest classifier is trained using default parameters and evaluated for each reference dataset using MLR [8].

### Hierarchical immune cell annotations

To integrate annotations and resolve differences across data sets, we developed a hierarchical set of cell types (Fig. 1b and Table S4). The cell types are found from Ontobee [9] (matching based on names and marker genes) and organized using the EBI Cell Ontology [10]. In this hierarchy, the top or root node represents the coarsest cell type, and successive levels describe increasingly finer ones. More precisely, child nodes are related to their parents via “is-a” relationships. This provides a framework to synthesize annotations from the reference data sets provided at different levels of granularity. Using this hierarchy, the 171 cell types (including redundancies) from the seven reference datasets are unified to a set of 35 non-redundant and hierarchical cell types (Fig. S2).

### Integrating annotations using a Deep Neural Network (DNN)

We predict the types in the hierarchy as a multi-class classification problem, leveraging the ability of DNNs to learn complex structure [11]. Cells were subsampled per cell type to avoid the dominance by cell number (Table S5). For each cell, the input layer takes the concatenation of the probability vectors from the dataset-specific classifiers and the output layer returns a vector of probabilities associated with hierarchical cell types (Fig. 1c). The position of each cell type in the hierarchy is encoded as a 35-bit vector that traces the path from the cell type to the root. There are three hidden layers with 200, 400 and 200 nodes, respectively, a topology that worked well in the training set, and the dropout rate is 0.2. The DNN is implemented as a multi-label multi-class classification model using keras [12] and tensorflow [13]. We used 10 trained DNN classifiers with identical hyper-parameters to produce a robust estimate of the mean and standard deviation of the probabilities for the subtypes. 10 DNN classifiers were trained using the balance dataset with epochs = 5 and batch_size = 4096.

### Hierarchical cell type assignment

For each query cell, we assigned its cell type based on the cell type hierarchy that exhibited the maximum entropy gain [14]. For each node (cell type) *c* on the cell type hierarchy, the entropy is calculated

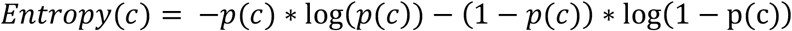

where p is the mean probability of that the query cell belongs to cell type *c* from the 10 DNN models above. Entropy is 0 when p equals to 0, by definition.

We adapted the well-known decision tree algorithm and revised the information gain from [15]

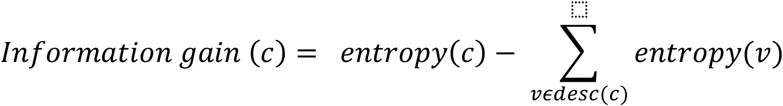

where *c* is the cell type and desc(*c*) is the set of child nodes of *c*.

This will select the cell type that can be assigned most confidently. To break ties in ranking, the cell type with the largest ratio of standard variation to mean is chosen.

### SingleR and Garnett predictions

SingleR annotation was generated by running SingleR using default parameters except for normalize.gene.length=T. Garnett annotation was generated by Garnett extended mode using the pre-trained classifier hsPBMC. The ImmClassifier annotation was generated using default parameters. Annotations were mapped across algorithms, as shown in Table S5.

### UMAP clustering and alluvial plots

The spatial coordinates of the cells were obtained using UMAP for each of the four datasets. The UMAP clustering of test datasets was performed using BETSY [16]. For dataset brca3p, brca5p and hcc, the top 500 most variable genes were used for PCA. Top 1000 most variable genes were chosen for dataset pbmc68k. Top 10 principle components, the nearest 50 neighbors and resolution=0.8 were used to generate the tSNE-plots. Alluvial plots were generated using ggalluvial [17].

