## Supplementary Materials for "Knowledge-based classification of fine-grained immune cell types in single-cell RNA-Seq data with ImmClassifier"

#### Supplementary Tables

**Table S1. A list of public scRNA-Seq and bulk-sequencing datasets used for evaluation datasets by ImmClassifier.**

| Dataset | Platform | Number of immune cells/samples | Number of immune cell types | Tissue,Cancer type/Normal | Ref |
|---|---|---|---|---|---|
| brca3p | inDrop,3' sequencing | 47,016 | 19 | PBMC, tumor, adjacent normal and lymph nodes, BRCA | [1] |
| brca5p | 10X, 5' sequencing | 28,013 | 8 | PBMC, tumor, adjacent normal and lymph nodes, BRCA | [1] |
| hcc | Smart-seq2 Full-length | 5,603 | 12 | PBMC, tumor and adjacent normal, HCC | [2] |
| hnscc | Smart-seq2 Full-length | 5,902 | 5 | Tumor and lymph nodes, HNSCC | [3] |
| pbmc68k | 10X, 3' sequencing | 68,579 | 11 | PBMC, Healthy | [4] |
| skcm | Smart-seq2 Full-length | 16,291 | 11 | Tumor, SKCM | [5] |
| bulk | Microarray | 211 | 38 | PBMC, BM, Healthy | [6] |

#### Table S2. Mapping IDs of cell types between different annotation methods.

| Dataset | Cell type | Original | ImmClassifier | SingleR | Garnett |
|---|---|---|---|---|---|
| Brca3p | Treg | T:reg | L:T:CD4:Treg | Treg* | |
| | CD4+T | T:CD4.* | L:T:CD4* | T_cells:CD4* | CD4 T cells |
| | CD8+T | T:CD8* | L:T:CD8* | T_cells:CD8* | CD8 T cells |
| | B | B | L:B | B_cell* | B cells |
| | NK | NK*(NKT excluded) | L:NK | NK* | NK cells |
| | mDC | mDC | M:cDC | DC:m* | Dendritic cells |
| | pDC | pDC | L:pDC,M:pDC | | |
| | MONOCYTE | MONOCYTE | M:Mono | Monocyte* | Monocytes |
| | MACROPHAGE | MACROPHAGE | M:Mac | Macrophage* | |
| | Neutrophil | NEUTROPHIL | M:Neu | Neutrophl* | |
| | Mast | MAST | M:Mast | | |
| | Unassigned | | | | Unknown |
| | T | | | | T cells |
| | NKT | NKT | | | |
| | Other | | CD34*, L:unconvT, M:Eos, M:Ery, M:Mega | Chondrocytes*, CMP, Embryonic_stem_cells, Endothelial_cells*, Erythroblast, Fibroblast*, GMP, HSC*, iPS_cells*,pre-B*, pro-B*, Smooth_muscle_cells*, Tissue_stem_cells* | |
| Brcap5p | T:CD4+Naive | T:CD4+Naive | L:T:CD4:Naïve | T_cell:CD4+_Naive | |
| | T:CD4+CM | T:CD4+CM | L:T:CD4:CM | T_cell:CD4+_central_memory | |

| | | | | | |
|---|---|---|---|---|---|
| | T:CD4+EM | T:CD4+EM | L:T:CD4:EM | T_cell:CD4+_effector_memory | |
| | T:CD8+Naive | T:CD8+Naive | L:T:CD8:Naïve | T_cell:CD8+_naive | |
| | T:CD8+CM | T:CD8+CM | L:T:CD8:CM | T_cell:CD8+_Central_memory | |
| | T:CD8+EM | T:CD8+EM | L:T:CD8:EM | T_cell:CD8+_effector_memory* | |
| | Treg | Treg | L:T:CD4:Treg | T_cell:Treg:Naive | |
| | CD4+T | | | T_cell:CD4+ | CD4 T cells |
| | CD8+T | | | T_cell:CD8+ | CD8 T cells |
| | T | | | | T cells |
| | Exhausted T | | L:T:CD4:Ex,L:T:CD8:Ex | T_cell:gamma-delta | |
| | Tissue-resident memory | | L:T:CD4:TRM,L:T:CD8:TRM | | |
| | Other | | CD34*,L:unconvT,M*,L:B*,L:T:CD4:Tfh, L:T:CD8:Mait | Tissue_stem_cells:*,B_cell*,Endothelial_cells*,Epithelial_cells*,Keratinocytes*,Macrophage*,Monocyte*,NK*,T_cell:gamma-delta | |
| hcc | cytotoxic CD4 cells | cytotoxic CD4 cells | | | |
| | effector memory CD8+T cells | effector memory CD8+T cells | L:T:CD8:EM | | |

| | | | | | |
|---|---|---|---|---|---|
| | exhausted CD4+ T cells | exhausted CD4+ T cells | L:T:CD8:Ex | | |
| | intermediate of EM and Exhausted T | intermediate of EM and Exhausted T | L:T:CD4:EM, L:T:CD8:EM, L:T:CD4:TRM, L:T:CD8:TRM, L:T:CD4:Ex, L:T:CD8:Ex | T_cell:effector_memory* | |
| | exhausted CD8+ T cells | exhausted CD8+ T cells | L:T:CD4:Ex | | |
| | MAIT cells | MAIT cells | L:T:CD8:Mait | | |
| | naive CD4+ T cells | naive CD4+ T cells | L:T:CD4:Naïve | T_cell:CD4+_Naive | |
| | Tregs | Tregs* | Treg | Treg* | |
| | naive CD8+ T cells | naive CD8+ T cells | L:T:CD8:Naïve | T_cell:CD8+_Naive | |
| | CD4+T | | | T_cell:CD4+ | |
| | CD8+T | | | T_cell:CD8+ | |
| | T helper | T helper | L:T:CD4:Tfh | | |
| | T | | | | T cells |
| | unknown | unknown | | | |
| | central memory T | | L:T:CD4:CM,L:T:CD8:CM | T_cell:Central_memory* | |

| | | | | | |
|---|---|---|---|---|---|
| | Other | | CD34, L:B*, L:NK, M* | Endothelial_cells*, Epitheial_cells, NK*,T_cell:gamma-delta | B cells, CD34+, Dendritic cells, Monocytes, NK cells |
| | Unassigned | | | | Unknown |
| pbmc68k | CD14+ Monocyte | CD14+ Monocyte | M:Mono | Monocyte | Monocytes |
| | CD19+ B | CD19+ B | L:B* | B_cell*, Plasma_cell* | B cells |
| | CD34+ | CD34+ | CD34* | CMP,GMP,MEP,CD34* | CD34 |
| | CD4+ T Helper2 | CD4+ T Helper2 | L:T:CD4:Tfh | | |
| | CD4+/CD45RA+/CD24- Naive T | CD4+/CD45RA+/CD24- Naive T | L:T:CD4:Naïve | T_cell:CD4+_Naive | |
| | CD4+/CD45RO+ Memory | CD4+/CD45RO+ Memory | L:T:CD4:CM,L:T:CD4:EM | T_cell:CD4+_central_memory,T_cell:CD4+_effector_memory | |
| | CD56+ NK | CD56+ NK | L:T:NK | NK* | NK cells |
| | CD8+ Cytotoxic T | CD8+ Cytotoxic T | | | |
| | CD8+/CD45RA+ Naive Cytotoxic | CD8+/CD45RA+ Naive Cytotoxic | L:T:CD8:Naïve | T_cell:CD8+_Naive | |
| | Dendritic | Dendritic | M:cDC,L:pDC,M:pDC | DC* | Dendritic cells |

| | | | | | |
|---|---|---|---|---|---|
| | CD4+T | | | T_cell:CD4+ | T_cell:CD4+ |
| | CD8+T | | | T_cell:CD8+ | T_cell:CD8+ |
| | CD8+ memory | | L:T:CD8:CM,L:T:CD8:EM | T_cell:CD8+_Central_memory,T_cell:CD8+_effector_memory* | |
| | CD4+/CD25 T Reg | | L:T:CD4:Treg | Treg* | |
| | T | | | | T cells |
| | Other | | L:T:CD4:TRM,L:T:CD8:TRM,M:Eos,M:Ery,M:Mac,M:Mega*,M:Neu,L:T:CD8:Mait, L:T:CD4:Ex, L:T:CD8:Ex | Platelets, pre-B*, pro-B*, T_cell:gamma-delta | |
| | unassigned | | | | Unknown |

**Table S3. A list of public scRNA-Seq datasets used as reference datasets by ImmClassifier.**

| Dataset | Platform | Number of immune cells/samples | Number of immune cell types | Tissue,Cancer type/Normal | Ref |
|---|---|---|---|---|---|
| hca-bm | 10X, 3' sequencing | 101,935 | 32 | Bone marrow, Normal | [7] |
| pbmc | 10X, 3' sequencing | 69,745 | 31 | PBMC, GI cancer | [8] |
| liver-immune | 10X, 3' sequencing | 4,906 | 6 | Liver, Normal | [9] |
| jci-bm | 10X, 3' sequencing | 73,246 | 18 | Bone marrow, Normal | [10] |
| nsclc-zilionis-tii-minor | inDrop, full-length | 34,558 | 34 | Tumor infiltrating immune cells, NSCLC | [9] |
| brcatil | 10X, 3' sequencing | 6,311 | 9 | Tumor infiltrating immune cells, TNBC | [11] |
| nsclc-guo | Smart-seq2 Full-length | 12,346 | 16 | Tumor infiltrating immune cells, NSCLC | [12] |

**Table S4. Full name of cell types in cell type hierarchy.**

| Full Name | Short Name |
|---|---|
| CD34 | CD34-positive stem cell |
| L | lymphocyte |
| M | myeloid cell |
| HSC | hematopoietic stem cell |
| pDC | plasmacytoid dendritic cell |
| cDC | conventional dendritic cell |
| Mast | mast cell |
| Mega | megakaryocyte |
| Ery | erythrocyte |
| Neu | neutrophil |
| Mono | monocyte |
| Mac | macrophage |
| Eos | eosinophil |
| T | T cell |
| NK | natural killer cell |
| B | B cell |
| unconvT | unconventional T cell |
| CD4+ | CD4-positive T cell |
| CD8+ | CD8-positive T cell |
| CD4+Tn | CD4-positive naïve T cell |
| CD4+Tcm | CD4-positive central memory T cell |
| CD4+Tem | CD4-positive effector memory T cell |
| CD4+Trm | CD4-positive tissue-resident memory T cell |
| CD4+Tex | CD4-positive exhausted T cell |
| Treg | regulatory T cell |

| | |
|---|---|
| Tfh | helper T cell |
| CD8+Tn | CD8-positive naïve T cell |
| CD8+Tcm | CD8-positive central memory T cell |
| CD8+Tem | CD8-positive effector memory T cell |
| CD8+Tex | CD8-positive exhausted T cell |
| CD8+Trm | CD8-positive tissue-resident memory T cell |
| MAIT | mucosal invariant T cell |
| PC | plasma cell |

**Table S5. The number of cells sampled to train deep neural network.**

| Cell number (N) | Sampled cell number per cell type |
|---|---|
| N<1000 | 1000 |
| 1000 <= N <2000 | 1200 |
| 2000 <= N <5000 | 1400 |
| N>=5000 | 1600 |

#### Supplementary Figures

### Figure S1

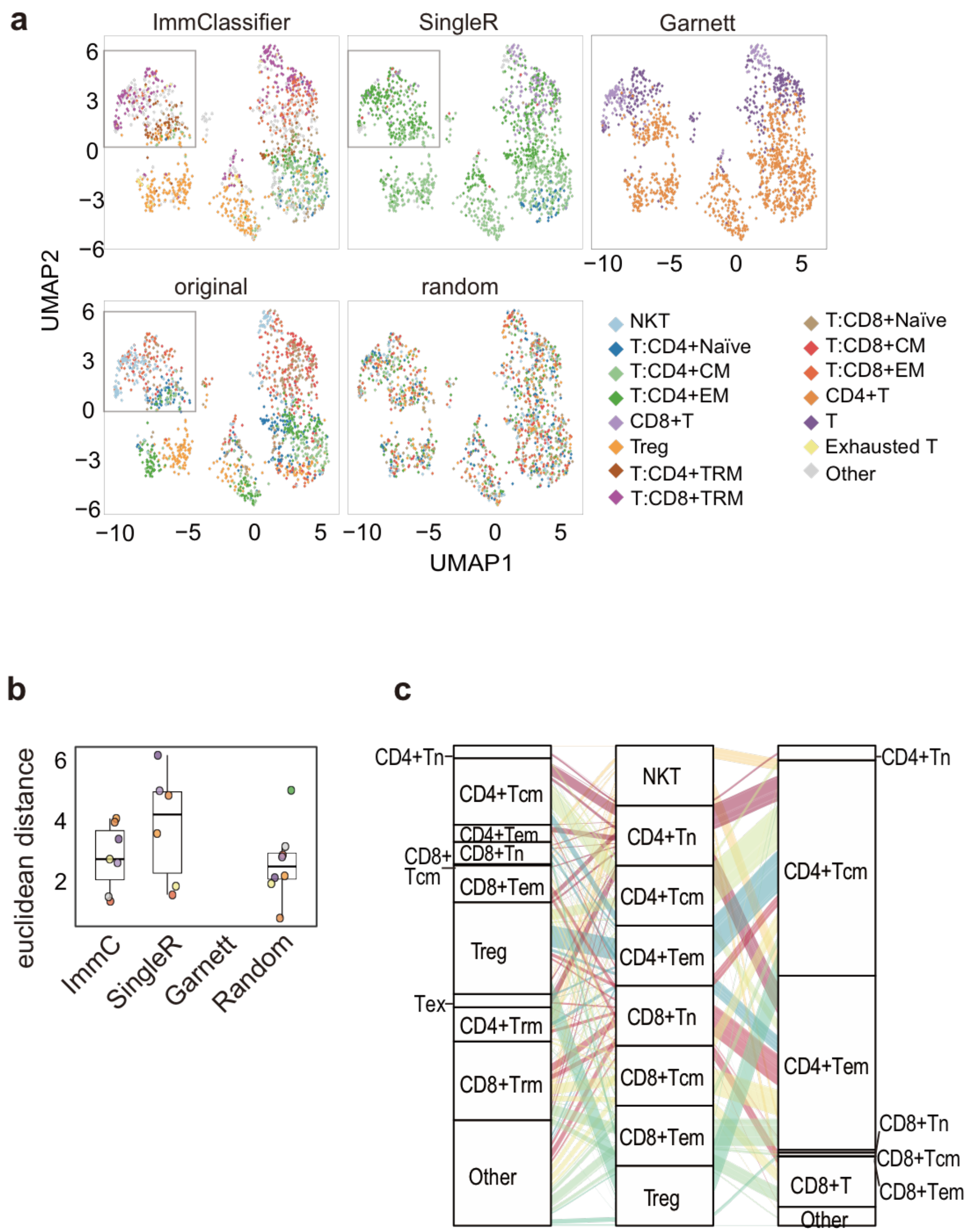

**Figure S1. Visualization of brca5p dataset by different annotation methods.** For better visualization, abundant cell populations (cell number >200) were sub-sampled to 200 per cell type. The mapping IDs of cell types between different annotations is listed in Table S5. If the annotation methods output a finer annotation granularity than the original annotation, the annotation terms of equivalent granularity to the original annotation were used in this figure. **(a)** UMAP plots colored by different annotation methods. In the first row (left to right), the panels are colored by ImmClassifier, SingleR and Garnett extended mode. In the second row (left to right), panels are colored by original publication, random shuffled cell labels. **(b)** Boxplots of Euclidean distance between centroids of predicted annotations and original annotations. (**c**) Alluvial plots connecting original annotated cell types to cell types annotated by ImmClassifier and SingleR.

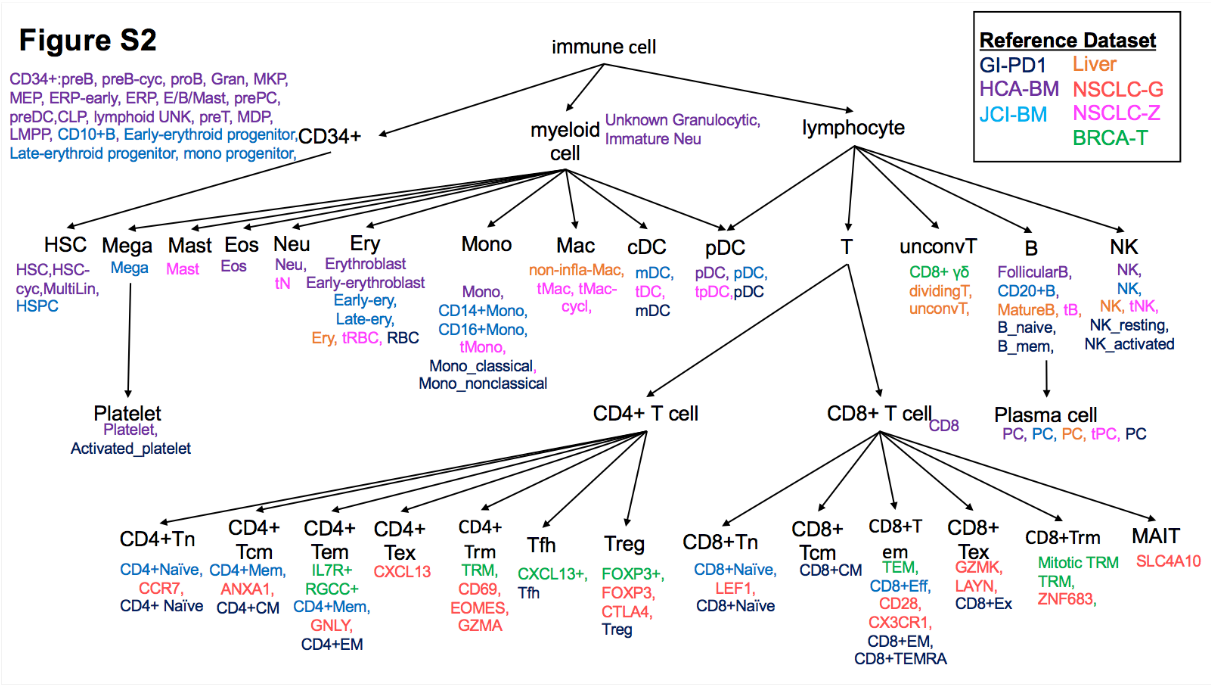


**Figure S2. Cell ontology hierarchy used by ImmClassifier.** The hierarchy is revised from cell differentiation hierarchy to accommodate for prediction probability. The hierarchy represents cell annotation granularity. Each node on the tree is a flat cell type. For each flat cell type (tree node), the matched cell types from training datasets are listed.

#### References (Supplementary Materials)


1. Azizi, E., et al., *Single-cell map of diverse immune phenotypes in the breast tumor microenvironment.* Cell, 2018. **174**(5): p. 1293-1308. e36.
2. Zheng, C., et al., *Landscape of infiltrating T cells in liver cancer revealed by single-cell sequencing.* Cell, 2017. **169**(7): p. 1342-1356. e16.
3. Puram, S.V., et al., *Single-cell transcriptomic analysis of primary and metastatic tumor ecosystems in head and neck cancer.* Cell, 2017. **171**(7): p. 1611-1624. e24.
4. Zheng, G.X., et al., *Massively parallel digital transcriptional profiling of single cells.* Nature communications, 2017. **8**: p. 14049.
5. Sade-Feldman, M., et al., *Defining T cell states associated with response to checkpoint immunotherapy in melanoma.* Cell, 2018. **175**(4): p. 998-1013. e20.
6. Novershtern, N., et al., *Densely interconnected transcriptional circuits control cell states in human hematopoiesis.* Cell, 2011. **144**(2): p. 296-309.
7. Hay, S.B., et al., *The Human Cell Atlas bone marrow single-cell interactive web portal.* Experimental hematology, 2018. **68**: p. 51-61.
8. Griffiths, J.I., et al., *Circulating immune cell phenotype dynamics reflect the strength of tumor-immune cell interactions in patients during immunotherapy.* Under review.
9. Zilionis, R., et al., *Single-cell transcriptomics of human and mouse lung cancers reveals conserved myeloid populations across individuals and species.* Immunity, 2019. **50**(5): p. 1317-1334. e10.
10. Oetjen, K.A., et al., *Human bone marrow assessment by single-cell RNA sequencing, mass cytometry, and flow cytometry.* JCI insight, 2018. **3**(23).
11. Savas, P., et al., *Single-cell profiling of breast cancer T cells reveals a tissue-resident memory subset associated with improved prognosis.* Nature medicine, 2018. **24**(7): p. 986.
12. Guo, X., et al., *Global characterization of T cells in non-small-cell lung cancer by single-cell sequencing.* Nature medicine, 2018. **24**(7): p. 978.
